# EXO70A2 is critical for the exocyst complex function in Arabidopsis pollen

**DOI:** 10.1101/831875

**Authors:** Vedrana Marković, Fatima Cvrčková, Martin Potocký, Přemysl Pejchar, Eva Kollárová, Ivan Kulich, Lukáš Synek, Viktor Žárský

## Abstract

Pollen development, pollen grain germination and pollen tube elongation are crucial biological processes in angiosperm plants that need precise regulation to deliver sperm cells to fertilize ovules. Pollen grains undergo two major developmental switches: dehydration characterized by metabolic quiescent state, and rehydration upon pollination that leads to extraordinary metabolic and membrane trafficking activity, resulting in germination and rapid tip growth of pollen tubes. To sustain these processes, many plant housekeeping genes evolved their pollen-specific paralogs. Highly polarized secretion at a growing pollen tube tip requires the exocyst tethering complex responsible for specific targeting of secretory vesicles to the plasma membrane. Here, we describe that EXO70A2 (At5g52340) is the main exocyst EXO70 isoform in Arabidopsis pollen, which governs the conventional secretory function of the exocyst, analogically to EXO70A1 (At5g03540) in the sporophyte. Our analysis of a CRISPR-generated *exo70a2* mutant revealed that EXO70A2 is essential for efficient pollen maturation, pollen grain germination and pollen tube growth. GFP:EXO70A2 was localized similarly to other exocyst subunits to the apical domain in growing pollen tube tips characterized by intensive exocytosis. Moreover, EXO70A2 could substitute for the EXO70A1 function in the sporophyte, indicating functional redundancy of these two closely related isoforms. Phylogenetic analysis revealed that the ancient duplication of EXO70A to two (or more) paralogs, one of which is highly expressed in pollen, occurred independently in monocots and dicots. In summary, EXO70A2 is a crucial component of the exocyst complex in the Arabidopsis pollen required for efficient plant sexual reproduction.

## INTRODUCTION

The pollen tube in Angiosperms represents an extremely elongated cellular structure, that emerges from a rehydrated pollen grain upon its landing on a stigma and delivers two sperm cells to fertilize an ovule. To reach the ovule, the pollen tube navigates through pistil tissues by highly polarized tip growth, restricted to the pollen tube apex. In this tightly regulated process, cell polarity maintenance mechanisms, addition of new membrane material and secretion of cell wall components are essential. Intensive exocytosis is localized to the very tip, followed by a subapical domain where endocytosis takes place to recycle the excess of membranes delivered in secretory vesicles (for review, see e.g. Hepler and Winship, 2015). Other key factors essential for the pollen tube tip growth include small GTPases and a Ca^2+^-signaling network regulating actin cytoskeleton dynamics (for review, see e.g. Cai et al., 2015). The cell wall of pollen tubes has a unique structure consisting of two layers: a pectinaceous and cellulose layer secreted at the apex, and an additional callose layer deposited in more distant regions from the tip. The spatial distribution of the cell wall components plays a critical role in the pollen tube morphogenesis (Chebli et al., 2012). Because pollen tubes are a simplified cell-autonomous system and exhibit highly polarized growth, they render an excellent model in cell biology for studies of cell polarity and regulation of secretion (e.g. Chebli et al., 2013; Qin and Dong, 2015). Precisely regulated membrane trafficking is crucial not only for pollen tube growth, but also for a previous stages of pollen development – pollen grain maturation and germination (Kang et al., 2003; Peng et al., 2011; Paul et al., 2016).

One of the fundamental regulators of polarized secretion is the exocyst – an evolutionarily conserved protein complex discovered due to its role in docking and tethering of secretory vesicles to specific sites at the plasma membrane (PM), facilitating subsequent fusion of secretory vesicles with the target membrane mediated by the SNARE proteins. The exocyst, first described in yeasts and mammals, consists of eight subunits – Sec3, Sec5, Sec6, Sec8, Sec10, Sec15, Exo70, and Exo84 (TerBush et al., 1996; Guo et al., 1999), with Sec3 and Exo70 generally believed to be responsible for targeting the complex to the PM, while the other subunits form the core of the complex (Boyd et al., 2004; He et al., 2007; Pleskot et al., 2015). Plant genomes encode all exocyst subunits (Cvrčková et al., 2001; Eliáš et al. 2003) that form a functional complex (Hála et al., 2008; Fendrych et al., 2010) and engage in cellular processes requiring targeted secretion, including root hair and pollen tube elongation (Cole et al., 2005; Synek et al., 2006; Hála et al., 2008; Synek et al., 2017), cytokinesis (Fendrych et al., 2010; Rybak et al., 2014), secondary cell wall deposition in trichomes and during xylem development (Kulich et al., 2015; Kubátová et al., 2019; Vukašinović et al., 2017), localized deposition of seed coat pectins (Kulich et al., 2010), transport of PIN auxin carriers to the PM (Drdová et al., 2013), and response to pathogens (Pečenková et al., 2011; Sabol et al., 2017). Viable mutants of *A. thaliana* defective in exocyst subunits (*sec8-4, sec15b, exo70a1, exo84b*) exhibit dwarfish growth with pleiotropic developmental defects (Cole et al., 2005; Synek et al., 2006; Hála et al., 2008; Fendrych et al., 2010; Batystová et al., submitted).

While in yeast and animals the exocyst subunits are typically encoded by a single gene, in land plants they are usually duplicated or even multiplicated (Eliáš et al., 2003; Cvrčková et al., 2012), with extreme gene proliferation in the case of the EXO70 subunit. For example, the genome of *Physcomitrella patens* encodes 13 *EXO70* paralogs, *Oryza sativa* 47 paralogs and *Arabidopsis thaliana* 23 paralogs, which can be divided into three clades of ancestral land plant origin, termed EXO70.1, EXO70.2 and EXO70.3 (Cvrčková et al., 2012; Rawat et al., 2017; Žárský et al., 2019). This plant-specific multiplicity of EXO70 suggests not only functional specialization in different plant tissues or cell types, but also a presence of several variants of the exocyst complex in the same cell (Žárský et al., 2013). Indeed, some EXO70.2 and EXO70.3 clade members contribute to conventional secretion in specific cell types, while several other isoforms acquired functions outside the conventional exocytosis pathway, acting in exocyst subcomplexes or even independently of the exocyst complex in some cases (Kulich et al., 2013; Kulich et al., 2015; Zhang et al., 2015; Hong et al., 2016; Pečenková et al., 2017; Synek et al., 2017). On the contrary, the Arabidopsis EXO70.1 isoform EXO70A1, highly expressed in most sporophytic tissues, is the main housekeeping EXO70 subunit participating in conventional exocytosis (Synek et al., 2006; Fendrych et al., 2010; Drdová et al., 2013). However, since EXO70A1 is not expressed in pollen, the main EXO70.1 paralog functioning in pollen remained experimentally uncharacterized until now.

The importance of the exocyst for pollen tube germination and growth is well documented. Homozygous Arabidopsis mutants in *SEC5a/b, SEC6, SEC8*, and *SEC15a* typically produce short and wide pollen tubes with compromised polarity, resulting in a male-specific transmission defect (Cole et al., 2005; Hála et al., 2008), and *sec3a* mutants cannot produce pollen tubes at all (Bloch et al., 2016). Transcriptomic and proteomic analyses documented high transcription of the *EXO70A2, C1, C2, F1, H3, H5*, and *H6* paralogs in Arabidopsis pollen, while EXO70C2, C1, and A2 were the most abundant in Arabidopsis pollen proteome (Grobei et al., 2009, Synek et al., 2017). The two closely related EXO70C paralogs from the EXO70.2 clade participate in the control of optimal pollen tube tip growth, probably via its regulatory function outside the exocyst complex (Synek et al., 2017). On the other hand, the clade EXO70.1 member EXO70A2 is the evolutionarily closest paralog to EXO70A1, the main sporophytic housekeeping isoform (Cvrčková et al., 2012), and also interacts with the same exocyst subunits as EXO70A1 (Synek et al., 2017). Therefore, EXO70A2 represents the best candidate for the main conventional EXO70 isoform in pollen functioning as a part of the exocyst complex in the regulation of polarized exocytosis.

In this report, we document that disruption of EXO70A2 in Arabidopsis affects pollen maturation, significantly reduces pollen germination efficiency and causes a severe pollen-specific transmission defect of the mutant allele due to impaired pollen tube growth. In growing pollen tubes, GFP:EXO70A2 localizes, similarly to other exocyst subunits, to the very apical domain in the tip characterized by intensive exocytosis. Ectopic expression of EXO70A2 could substitute for the EXO70A1 function in the sporophyte, indicating functional redundancy of these two closely related isoforms. Phylogenetic and expression analysis of EXO70.1 genes revealed independent duplications of EXO70A in monocots and dicots and showed that the dicot EXO70A2 clade contains predominantly pollen-specific isoforms. We thus conclude that Arabidopsis EXO70A2 is critical for the pollen function and that, analogous to EXO70A1 in the sporophyte, EXO70A2 is the main EXO70 isoform functioning as a subunit of the exocyst complex in conventional secretion in Arabidopsis pollen.

## RESULTS

### EXO70A2 is a member of a dicot pollen-expressed EXO70.1 clade

To clarify the evolutionary relationships among EXO70A paralogs, which comprise the sole family of the ancestral EXO70.1 clade (Synek et al. 2006; Cvrčková et al. 2012; Rawat et al. 2017), we performed a detailed phylogenetic analysis of available predicted protein sequences of this clade members from 19 plant species covering a broad range of land plant diversity including liverworts, mosses, lycophytes, gymnosperms, the basal angiosperm *Amborella trichopoda*, grasses as representatives of monocots, and dicots including representatives of both asterids and rosids (sequences are listed in Supplemental File S1). The sole EXO70 of the charophyte *Klebsormidium flaccidum* has been also included as an outgroup. The resulting phylogenetic tree (Fig. 1) clearly indicates that the EXO70A duplication took place independently in basal dicots and in monocots (or at least in grasses), resulting in both dicots and grasses possessing representatives of two EXO70A clades that cannot be considered orthologous between the two angiosperm lineages. The Arabidopsis EXO70A1 and EXO70A2 paralogs map into different dicot EXO70A clades, while the third, EXO70A3, is closely related to EXO70A2 and resulted from a recent gene duplication that may have been restricted to Brassicales, or even to some subgroup thereof.

**Figure 1.**
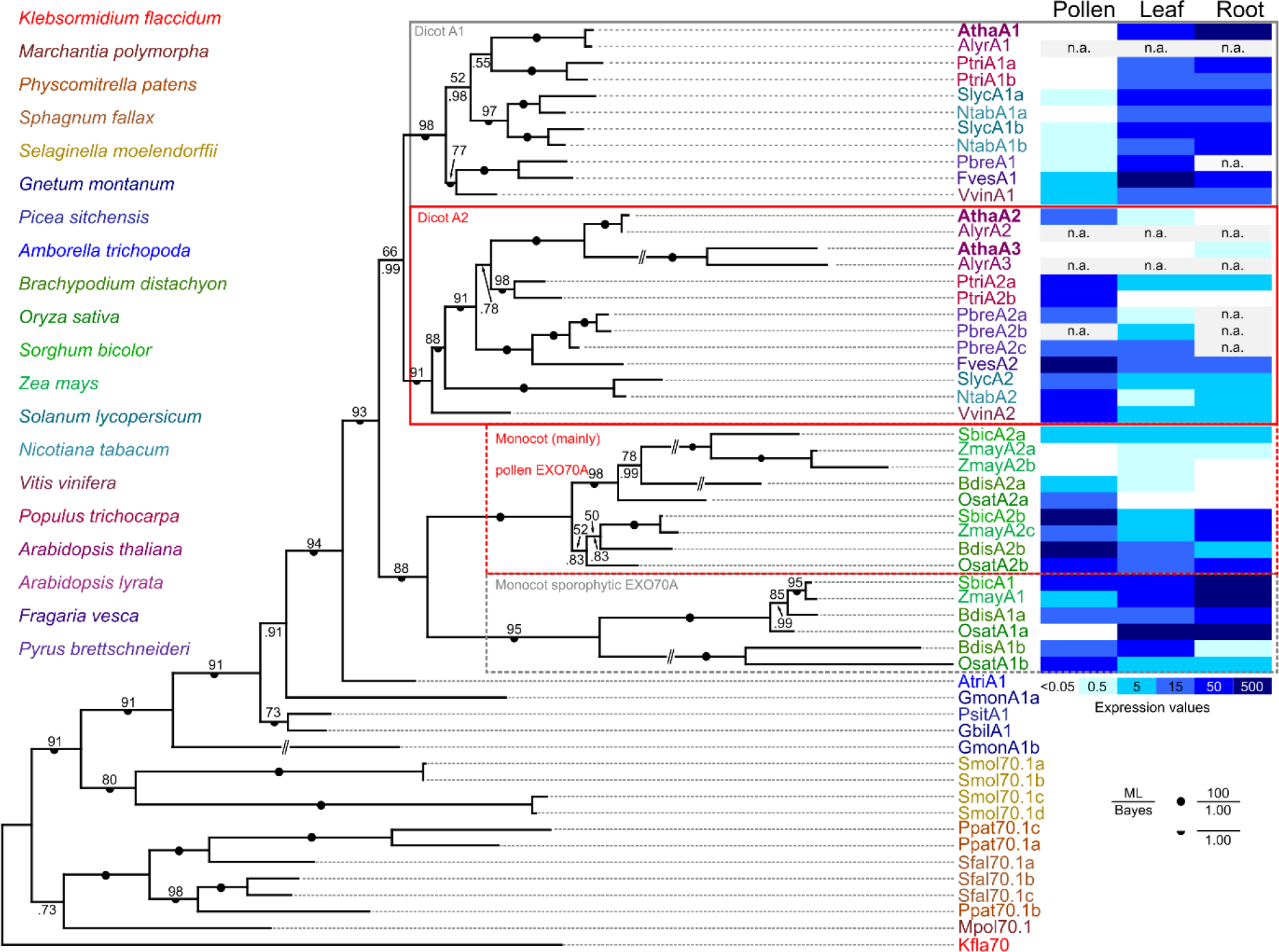
Phylogenetic and expression analysis of the EXO70.1 clade in land plants. A Bayesian phylogenetic tree of EXO70.1 protein sequences from representative land plants ranging from liverworts (*Marchantia polymorpha*), mosses (*Physcomitrella patens, Sphagnum fallax*), lycophytes (*Selaginella moelendorffii*), gymnosperms (*Gnetum sp., Picea sitchensis*), the basal angiosperm *Amborella trichopoda*, grasses (*Brachypodium distachyon, Oryza sativa, Sorghum bicolor* and *Zea mays*) as representatives of the monocots, and dicots represented by both asterids (*Solanum lycopersicum, Nicotiana tabacum*) as well as multiple rosids (*Vitis vinifera, Populus trichocarpa, Arabidopsis thaliana and lyrata, Fragaria vesca, Pyrus brettschneideri*). The sole EXO70 of the charophyte *Klebsormidium flaccidum* has been included as an outgroup (for a full list of sequences with accessions see Supplemental File S1). A consistent tree was also obtained by the maximum likelihood method (bootstrap values shown); full support for some branches is denoted by symbols. For selected angiosperm species, a heat map of transcript levels in pollen and sporophytic tissues, as inferred from publicly available transcriptome data, is shown.

For selected representatives of both monocots and dicots, transcript level data for the analyzed genes were extracted from public databases and mapped onto the phylogenetic tree (Fig. 1). The results document clade-specific distinct expression patterns in both dicots and grasses. In dicots, members of the clade containing Arabidopsis EXO70A1 usually exhibit high expression levels in sporophytic vegetative organs with low or no expression in pollen, while a complementary pattern, i.e. high transcript level in pollen and lower to no expression in the sporophyte, is typical for the clade containing EXO70A2. Also in the grasses one of the two clades contains mostly genes with either ubiquitous or predominantly sporophytic expression, while most members of the other exhibit enhanced, though usually not exclusive, expression in the male gametophyte.

### Preparation and characterization of a loss-of-function *exo70a2* allele using the CRISPR/Cas9 system

In order to investigate the *EXO70A2* function *in planta* we employed Arabidopsis mutants. At the time of this study, T-DNA insertion null mutants in *EXO70A2* (At5g52340) were not available. One of the two publicly available lines, GABI_824D06 (*exo70a2-1*) with the insertion located in 5’ UTR, exhibited EXO70A2 overexpression (Synek et al., 2017), and the another, FLAG_264F01 (*exo70a2-2*) could not be confirmed in our hands. We thus employed the CRISPR/Cas9 system (Wang et al., 2015) to generate a loss-of-function *exo70a2* mutant in the Arabidopsis Col-0 background, and obtained a mutant line (*exo70a2-3*) bearing a 13-bp insertion in the fifth exon of the *EXO70A2* gene, causing a frameshift in the CDS (Supplemental Fig. S1A). This mutation probably affected the *EXO70A2* RNA stability or processing, because the *EXO70A2* transcript level was reduced to less than half in the *exo70a2-3* mutant as evaluated by semi-quantitative RT-PCR in flower material (Supplemental Fig. S1B).

Plants homozygous for the CRISPR-generated insertion, hereafter referred to as *exo70a2* only, were then subjected to phenotype analysis and compared to their wild-type siblings. To address whether EXO70A2 might play some role in the sporophyte, despite its minimal expression in sporophytic tissues (Synek et al., 2006; www.Genevestigator.com – Hruz et al., 2008), we inspected the morphology of the mutant and WT plants and evaluated two basic morphological parameters. Primary root length of 7 days old seedlings grown on vertical agar was comparable between exo70a2 and WT (Supplemental Fig. S1C, D). Similarly, 5 weeks old *exo70a2* and WT plants showed no differences in the plant height and general plant morphology (Supplemental Fig. S1E, F). In conclusion, the sporophyte of *exo70a2* homozygotes was not affected by the mutation, consistent with EXO70A2 being a pollen-specific EXO70 isoform.

### The *exo70a2* mutant shows a pollen-specific transmission defect

We next analyzed the transmission of the *exo70a2* mutant allele through pollen. Based on PCR genotyping, the progeny of heterozygous *exo70a2* mutant plants showed significantly reduced portion of mutant homozygotes in comparison to the normal Mendelian ratio (only 15% of mutant homozygotes were detected), pointing to a severe transmission defect of this mutant allele (Tab. 1).

**Table 1.** Segregation Of The Heterozygous *exo70a2* Mutant.

| | +/+ | +/- | -/- | n | $\chi^2$ | P |
| --- | --- | --- | --- | --- | --- | --- |
| Expected | 25.0% | 50.0% | 25.0% |  |  |  |
| *exo70a2* | 38.3% | 46.7% | 15.0% | 107 | 12.14 | 0.002 |

+/+, wild-type plants; +/-, heterozygous mutants; -/-, homozygous mutants

To determine whether the transmission defect was male- or female-specific, we performed reciprocal crossing of *exo70a2* heterozygotes to WT plants and inspected the frequency of WT and heterozygous progeny. When *exo70a2* heterozygous plants were used as pollen donors, only 17% of the progeny were heterozygous for the *exo70a2* allele instead of expected 50%. However, when they were used as pollen recipients, 54% of the progeny was heterozygous, indicating that the transmission defect was male-specific (Tab. 2).

**Table 2.** Reciprocal crossing of the *exo70a2* mutant and wild type (Col-0).

| Pollen donor +/-; Pollen recipient, +/+ |  |  |  |  |  |
| --- | --- | --- | --- | --- | --- |
| | +/+ | +/- | n* | $\chi^2$ | P |
| Expected | 50.0% | 50.0% |  |  |  |
| *exo70a2* | 82.6% | 17.4% | 69 | 29.35 | <0.001 |

| Pollen donor +/+; Pollen recipient +/- |  |  |  |  |  |
| --- | --- | --- | --- | --- | --- |
| | +/+ | +/- | n* | $\chi^2$ | P |
| Expected | 50.0% | 50.0% |  |  |  |
| *exo70a2* | 46.2% | 54.8% | 93 | 0.53 | 0.47 |

+/+, wild-type plants; +/-, heterozygous mutants
\* at least 6 independent crossings were prepared

### Pollen maturation, germination and pollen tube growth are compromised in the *exo70a2* mutant

To investigate the nature of the pollen-specific transmission defect, we inspected several phases of pollen development, including pollen germination and pollen tube growth in *exo70a2* and WT plants.

We observed four times higher percentage of non-viable pollen grains at the mature pollen stage using Alexander staining, indicating a significant difference (WT: 3.4%, n = 179; *exo70a2*: 11.5%, n = 157; Chi-square test p_value = 0.004) (Supplemental Fig. S2A). However, DAPI staining of pollen nuclei demonstrated that *exo70a2* pollen grains could normally reach the tricellular developmental stage. This suggests that the observed defect occurred in the maturation phase (Supplemental Fig. S2B).

We then conducted *in vitro* germination assay on pollen samples harvested from homozygous and heterozygous *exo70a2* mutants as well as from a WT control. While the control pollen started to efficiently germinate within two hours after imbibition, the pollen from *exo70a2* homozygotes germinated not sooner than only eight hours after the imbibition (Fig. 2A, B). Even after 20 hours the *exo70a2* pollen still showed 8-fold lower germination efficiency than WT pollen. Since high concentrations of extracellular reactive oxygen species (ROS) were previously reported at the onset of the pollen germination process (Smirnova et al., 2014), we analyzed the ROS production in WT and *exo70a2* pollen grains 20 min after imbibition using histochemical staining of superoxide anion radicals by nitroblue tetrazolium (NBT). The ROS levels were dramatically reduced in mutant pollen grains, indicating their different physiological status (Fig. 2C, D).

**Figure 2.**
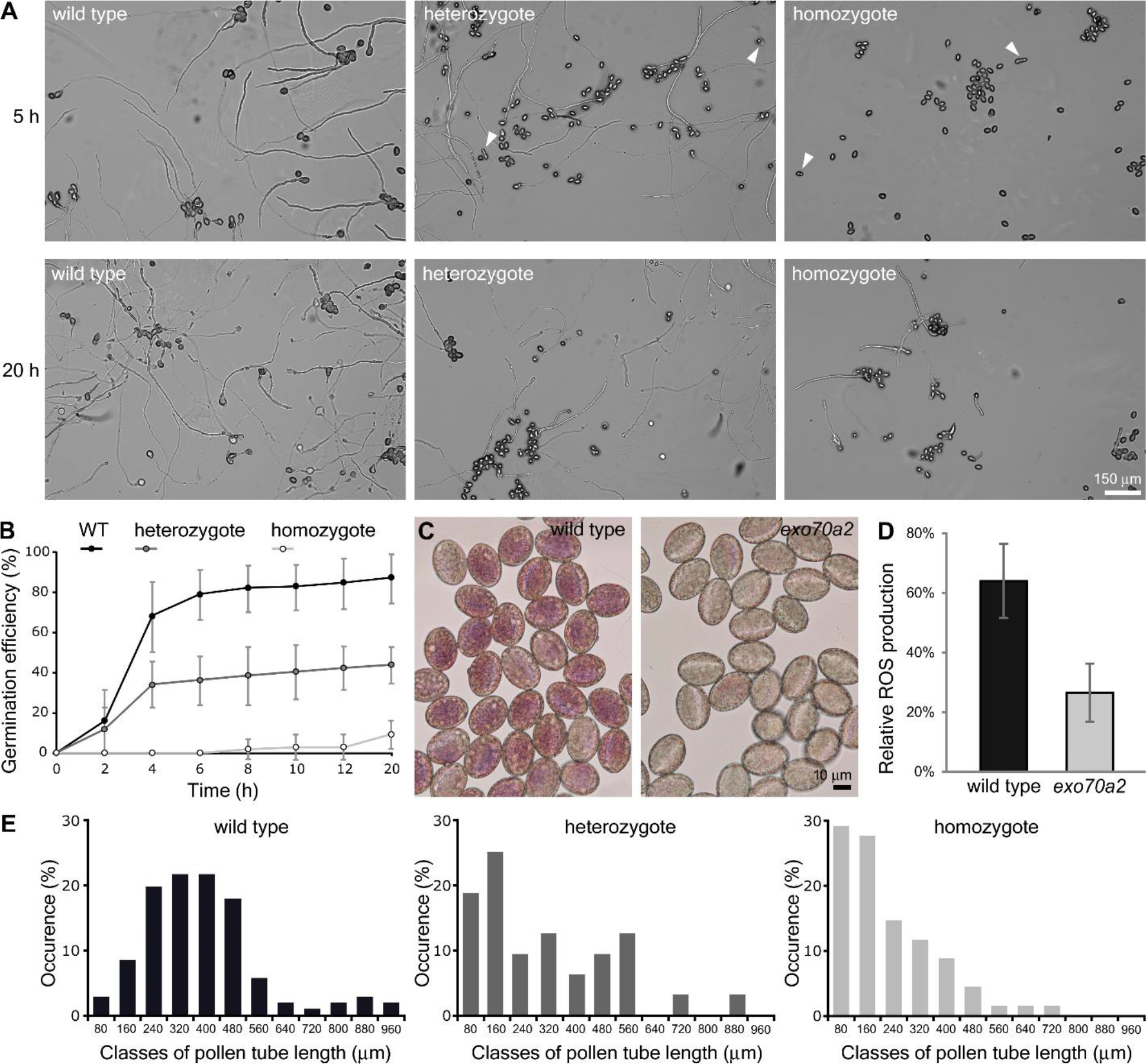
*in vitro* pollen germination and pollen tube growth defects of *exo70a2* mutants. A) Representative micrographs of *in vitro* germinated pollen from WT and heterozygous and homozygous EXO70A2 mutant plants 5 h and 20 h after imbibition. Arrowheads indicate germinated mutant pollen tubes. ) Pollen germination efficiency *in vitro* at different time points (10 samples originating from 3 different plants were evaluated for each genotype, error bars represent SD). C) ROS production 20 min after pollen imbibition as visualized by NBT staining. D) Quantification of the ROS staining (in C) shows a significant difference between WT and *exo70a2* pollen grains (error bars represent SD, t-test p_value < 0.001). E) Distribution of pollen tube lengths 20 h after imbibition. More than 100 pollen tubes were evaluated for WT and heterozygous plants, and 50 pollen tubes for the *exo70a2* mutant.

To find out whether the mutation only affects pollen tube emergence or also its subsequent elongation, we analyzed pollen tube lengths 20 hours after imbibition *in vitro*. Indeed, in this experiment, *exo70a2* pollen tubes were dramatically shorter than WT ones (Fig. 2A, E). Subsequent analysis of growth dynamics of individual pollen tubes revealed that the averaged maximal growth rate of *exo70a2* pollen tubes was 6 times lower in our conditions than that of their WT counterparts (*exo70a2*: 0.39 ± 0.08 μm-min^−1^, WT: 2.42 ± 0.37 μm-min^−1^, n > 12; mean ± SD; Student’s t-test p_value < 0.0001). Thus, *exo70a2* pollen displays both delayed germination and reduced pollen tube growth rate. Interestingly, an inspection of the *exo70a2* pollen tube elongation *in vivo* in self-pollinated pistils showed less dramatic, yet significant, defect in pollen tube growth when compared to WT (Fig. 3). This can be explained by the presence of chemical signals promoting pollen tube growth in a dose-dependent manner produced by the transmitting tract tissues in pistils (Johnson and Preuss 2002; Vogler et al., 2014).

**Figure 3.**
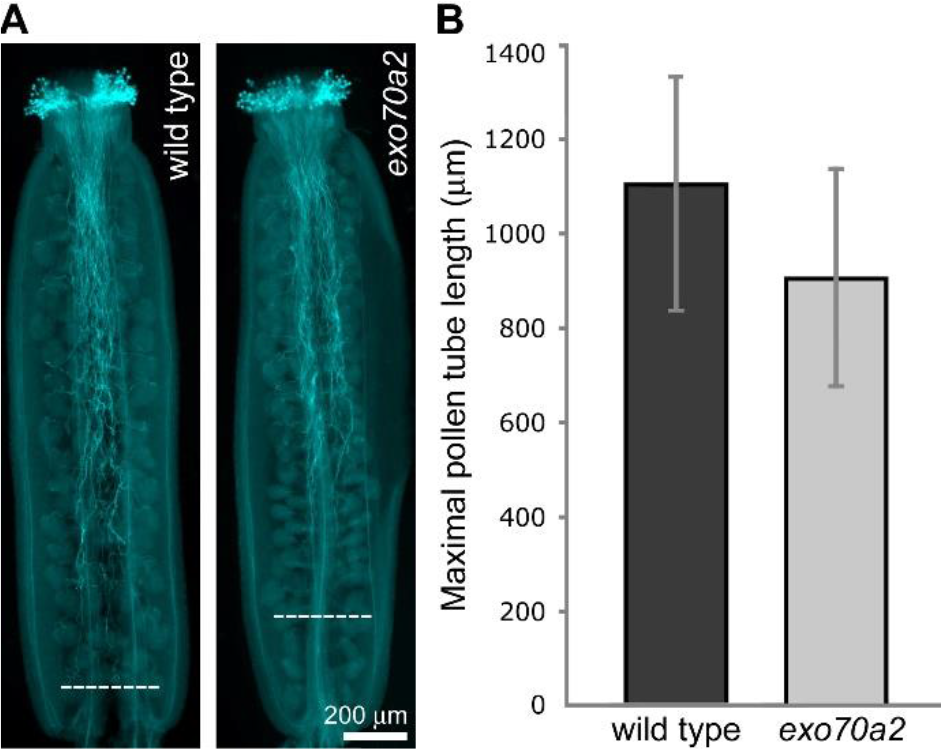
*In vivo* growth of *exo70a2* and WT pollen tubes. A) Aniline blue staining visualizing callose in pollen tubes in representative self-pollinated pistils from a WT and homozygous *exo70A2* plant. Dashed lines indicate the longest pollen tubes. B) Quantification of maximal pollen tube lengths based on aniline blue staining (at least 30 pistils of each genotype were evaluated, error bars represent SD, Student’s t-test p_value < 0.01).

Taken together, the loss of functional EXO70A2 affects not only pollen tube growth but also pollen maturation and germination. These processes are linked to polarized exocytosis, and similar defects have been observed in other exocyst mutants (Cole et al., 2005; Hála et al., 2008; Bloch et al., 2016).

### Slow growing *exo70a2* pollen tubes display morphological defects without major changes in the cell wall composition

In the *in vitro* germination experiments, we noticed that *exo70a2* pollen tubes had grown straight, without any branching, but with about 50% wider diameter than WT ones (Fig. 4). This morphological defect suggests that secretion is delocalized in the apex of *exo70a2* pollen tubes, which is consistent with the canonical exocyst function in polarized exocytosis in the tip growth.

**Figure 4.**
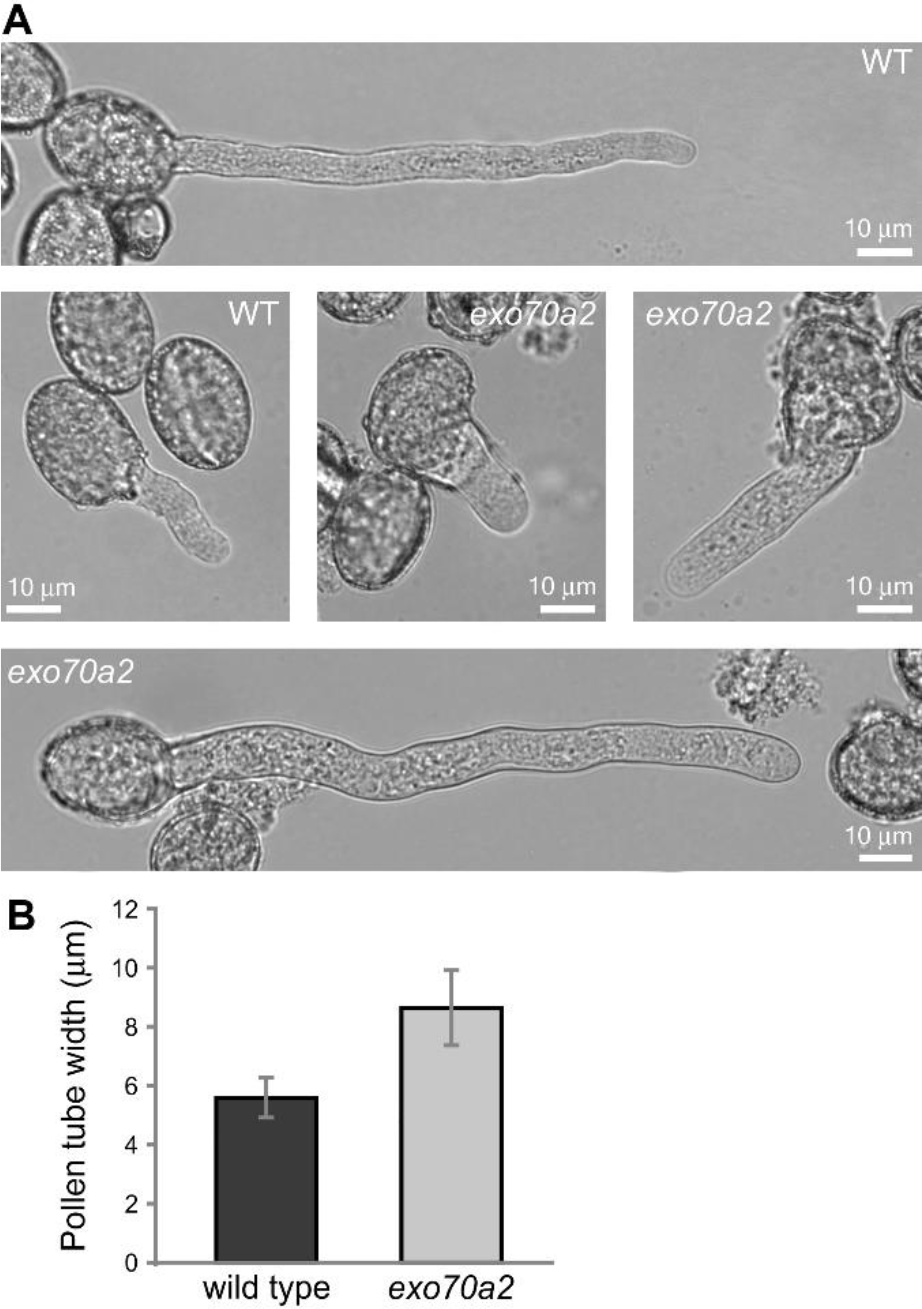
Morphology of *exo70a2* and WT pollen tubes *in vitro*. A) WT and *exo70a2* pollen tubes 1.5 h and 8h, respectively, after imbibition. B) Measurement of WT and *exo70a2* pollen tube width showed a significant difference (n > 46; Student’s t-test p_value < 0.0001).

Since the exocyst plays an important part in polarized exocytosis, which, in turn, is crucial for cell wall construction, we tested whether the pollen tube growth defect in *exo70a2* might be due to altered cell wall biogenesis. Therefore, we analyzed the distribution of dominant cell wall components in growing pollen tubes *in vitro*. Nevertheless, pectins showed normal distribution after propidium iodide staining, and cellulose and callose deposition visualized by calcofluor white and aniline blue staining, respectively, only extended more toward the tip in *exo70a2* pollen tubes, which could be explained by their much lower growth rate (Fig. 5).

**Figure 5.**
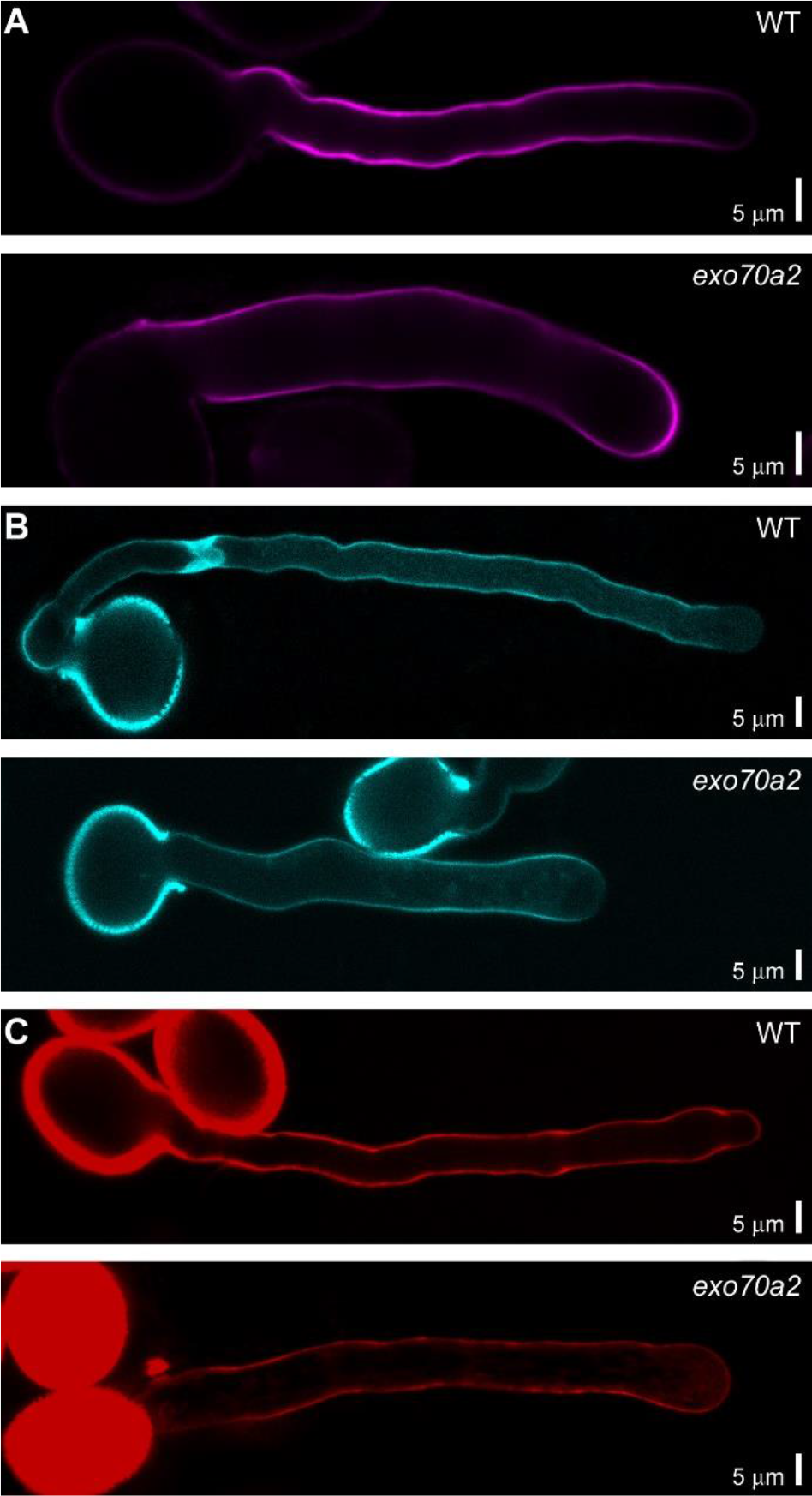
Cell wall deposition in growing WT and
*exo70a2* pollen tubes *in vitro*. A) Visualization of cellulose by calcofluor white staining. B) Visualization of callose by aniline blue staining. C) Visualization of pectins by propidium iodide staining.

### GFP:EXO70A2 complements the *exo70A2* mutation and localizes to the PM in pollen tube tips

For further functional characterization of EXO70A2, we prepared a GFP-tagged variant of *EXO70A2* expressed under its native promoter. The *pEXO70A2∷GFP:EXO70A2* construct was introduced into *exo70A2* heterozygous plants in order to analyze the EXO70A2 localization and functionality in mutant and WT pollen. The expression of *pEXO70A2∷GFP:EXO70A2* complemented the *EXO70A2* disruption: The percentage of non-viable pollen grains in freshly open flowers was similar to WT (complemented line: 3.9%, n = 231; WT: 3.4%, n = 179; Chi-square test p_value = 0.772). The ability of *exo70a2* pollen to germinate *in vitro* without any delay was restored (Fig. 6A, B; Supplemental Fig. S3), the pollen tube growth defect was reverted to normal (Fig. 6C) and also the pollen tube width was similar to WT tubes (mean ± SD; complemented line: 6.20 ± 0.44 μm; WT: 6.37 ± 0.72 μm, n > 32; Student’s test p_value < 0.2631). These observations confirm that the reported defects in *exo70a2* pollen were due to the disruption of the *EXO70A2* gene.

**Figure 6.**
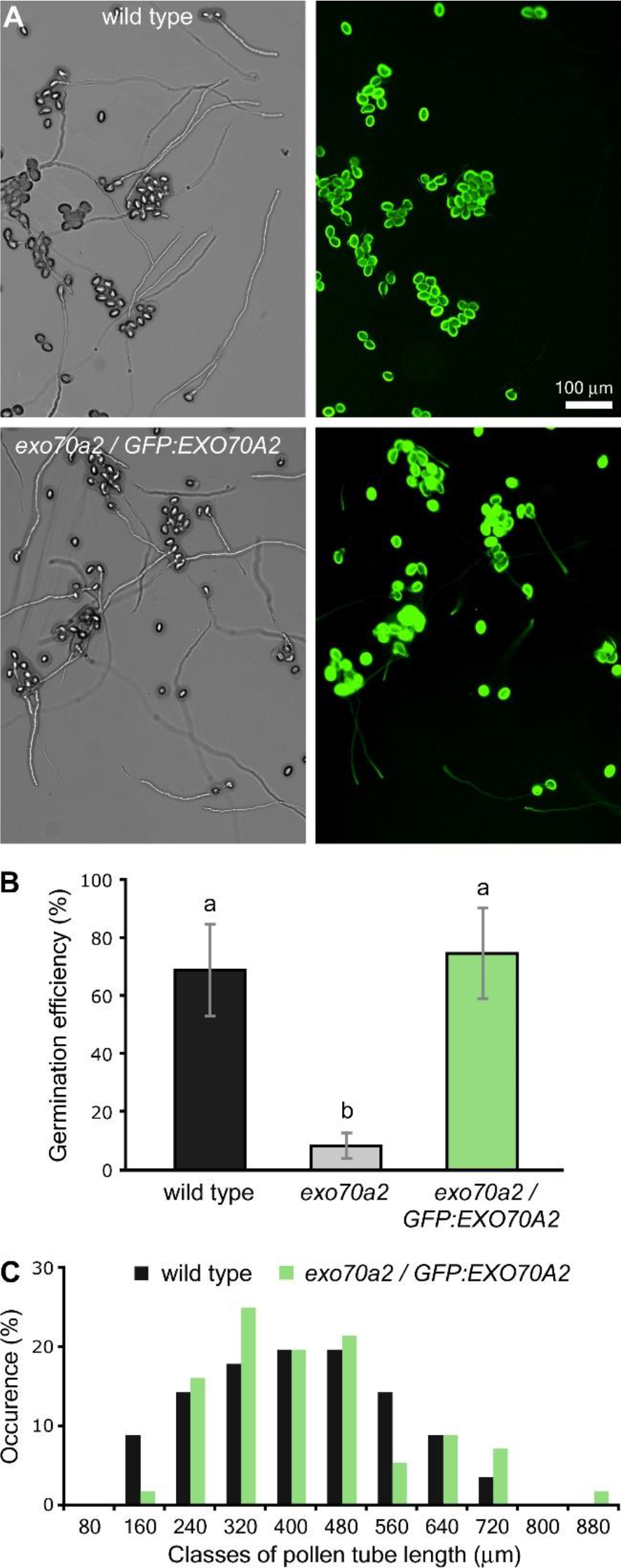
Expression of *pEXO70A2∷GFP:EXO70A2* complements the defects of the EXO70A2 mutant pollen. A) Representative micrographs of *in vitro* germinated pollen from WT and the *exo70a2/pEXO70A2∷GFP:EXO70A2* line 20 h after imbibition. B) Pollen germination efficiency *in vitro* 20 h after imbibition (10 samples originating from 3 different plants were evaluated for each genotype, error bars represent SD). Letters denote statistically different groups evaluated by ANOVA at 0.01 significance level. C) Distribution of pollen tube lengths 20 h after imbibition. More than 60 pollen tubes for each genotype were evaluated.

While seedlings and mature plant organs exhibited no fluorescence signal of GFP:EXO70A2 (not shown), we clearly detected GFP:EXO70A2 in pollen grains and pollen tube tips (Fig. 7). In the unicellular pollen stage, GFP:EXO70A2 was found mostly in the nucleus; in the bicellular pollen stage, it was partially re-localized to the cytoplasm; and in the later stages, it was evenly distributed throughout the cytoplasm, but completely excluded from nuclei (Fig. 7A). Importantly, in complemented *exo70a2* pollen tubes, GFP:*EXO70A2* displayed a highly polarized localization along the PM in the apical domain of growing pollen tube tips with a minor portion in the cytoplasm (Fig. 7B), which is a typical pattern for exocyst subunits (Synek et al., 2017). In the WT background, however, the polar PM localization was absent and GFP:EXO70A2 was distributed entirely in the cytoplasm of pollen tubes (Fig. 7C), which could be explained by a competition of the native and GFP-tagged *EXO70A2* variants for binding into the exocyst complex as observed previously in tobacco pollen tubes (Sekereš et al., 2017), and also in mammalian cells (Matern et al. 2001).

**Figure 7.**
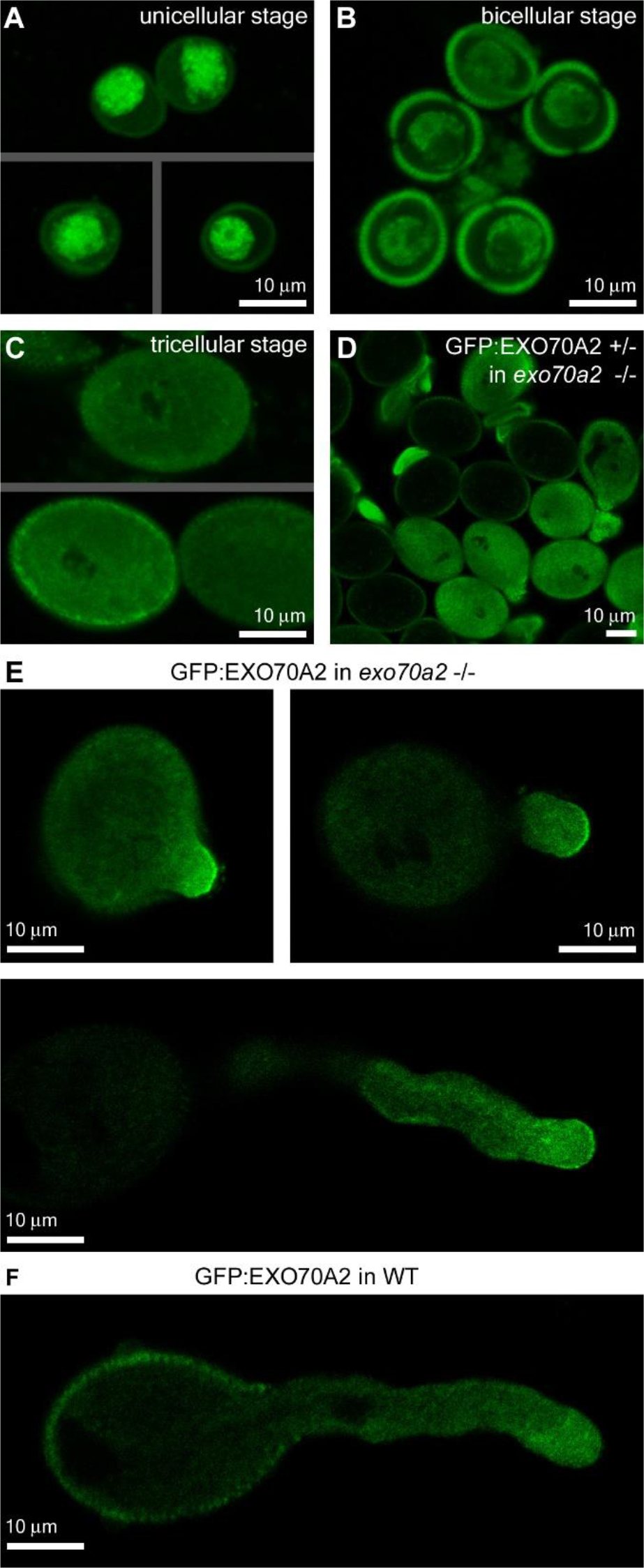
Localization of GFP:EXO70A2 expressed under its native promoter in pollen. A) GFP:EXO70A2 localization in unicellular pollen grains in the *exo70a2* background. B) GFP:EXO70A2 localization in bicellular pollen grains in the *exo70a2* background. C) GFP:EXO70A2 localization in tricellular pollen grains in the *exo70a2* background. Note that images in A, B and C were captured at an identical setting, allowing for their quantitative comparison. D) GFP:EXO70A2 localization in mature *exo70a2* pollen grains in a line segregating for the *GFP:EXO70A2* cassette, providing an internal control for pollen grain autofluorescence. E) GFP:EXO70A2 localization in different stages of pollen tube elongation in the *exo70a2* background. F) GFP:EXO70A2 localization in a pollen tube in the WT background

### EXO70A2 can rescue the EXO70A1 disruption in the sporophyte

We were interested what is the degree of functional specialization versus redundancy of EXO70A1 and EXO70A2 paralogs implied by their close sequence similarity (see Fig. 1). In order to address it experimentally we expressed *EXO70A2* and *EXO70A1*, respectively, under the control of the *EXO70A1* promoter in the *exo70a1* mutant background and analyzed EXO70A2 localization and functionality in the sporophyte. The *exo70a1* mutant was earlier shown to exhibit pleiotropic morphological defects, including retarded polar growth of root hairs, a loss of apical dominance, dwarfish stature and sterility (Synek et al., 2006). Moreover, GFP:EXO70A1 fully complemented the *exo70a1* loss-of-function allele and localized at the PM, with special enrichment at the outer lateral domain in root epidermal cells (Drdová et al., 2013; Fendrych et al., 2013).

We found that the expression of *pEXO70A1∷GFP:EXO70A2* rescued the morphological defects of *exo70a1* mutant plants similarly to *pEXO70A1∷GFP:EXO70A1*, including plant height of 7-week-old plants (Fig. 8A, B). Moreover, the GFP:EXO70A2 localization was identical to GFP:EXO70A1 in the root epidermal cells, suggesting that EXO70A1 and EXO70A2 paralogs are functionally redundant in the sporophyte (Fig. 8C).

**Figure 8.**
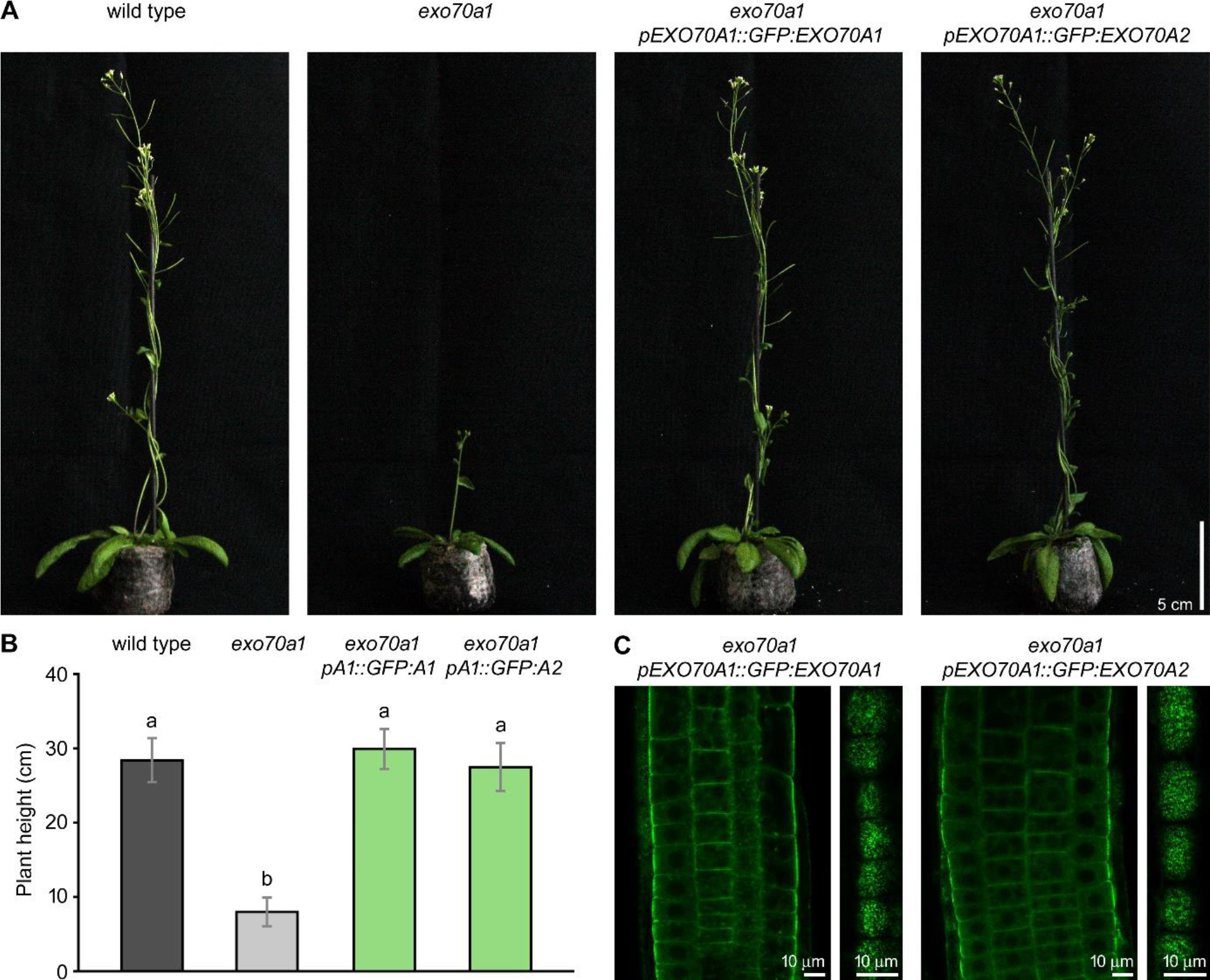
Ectopic expression of EXO70A2 in sporophytic tissues of *exo70a1* mutant plants. A) Representative images of 40-day-old plants document that the *pEXO70A1∷GFP:EXO70A2* expression in *exo70a1* mutant plants rescued its growth defects similarly to *pEXO70A1∷GFP:EXO70A1*. B) Measurement of 40-day-old plant height. Bars represent SD; letters denote statistically different groups calculated by ANOVA at 0.01 significance level. C) *pEXO70A1∷GFP:EXO70A2* shows very similar localization pattern to *pEXO70A1∷GFP:EXO70A1* in root epidermal cells. Left panels - confocal sections through the root transition zone. Right panels - confocal sections in the level of lateral plasma membranes.

## DISCUSSION

In order to deliver sperm cells to ovules and achieve successful fertilization, highly efficient secretory machinery is essential to sustain the fast pollen tube growth. Targeting of secretory vesicles to a very small PM area in the growing pollen tube tip requires, among other regulators, also the vesicle-tethering exocyst complex. The crucial role of the exocyst in the tip growth was previously evidenced by the aberrant morphology of pollen tubes in Arabidopsis mutants deficient in core exocyst subunits (Cole et al., 2005; Hála et al., 2008). However, the main EXO70.1 isoform contributing to the conventional secretory function of the exocyst complex in pollen awaited experimental characterization until now.

We previously described that EXO70.2 clade members, EXO70C2 and EXO70C1, are not stable subunits of the exocyst complex (Synek et al., 2017), leaving EXO70A2 the best candidate for the main housekeeping EXO70 isoform in pollen. First, EXO70A2 is the closest paralog to EXO70A1 – the main EXO70 isoform in the Arabidopsis sporophyte – and these two share 72% sequence identity at the protein level. Second, EXO70A2 also physically interacted with three other exocyst subunits (SEC3a, SEC10b and EXO84b), similarly to EXO70A1 (Hála et al., 2008; Fendrych et al., 2010; Synek et al., 2017). Third, the expression patterns of *EXO70A1* and *EXO70A2* alternate between the sporophyte-specific and pollen-specific, respectively. This is in agreement with the hypothesis suggesting that after a whole-genome duplication in plants, genes directly linked to cell polarity tend to diverge in their expression patterns to sporophyte-specific and to those enriched in tip-growing cells, especially in the male gametophyte (De Smet et al., 2017). In general, about 10% of essential housekeeping genes evolved their pollen specific paralogs to sustain the specific requirements for tip growth (Becker and Feijó, 2007).

Indeed, in course of the extraordinary diversification and functional specialization of the large EXO70 family in angiosperms (Cvrčková et al., 2012), the EXO70A duplication to sporophyte- and male gametophyte-specific paralogs occurred independently in monocots and dicots. In some plant taxonomic groups, each of them further duplicated, generating, for example, EXO70A2 and EXO70A3 in Brassicaceae, and EXO70C1a and EXO70C1b in Solanaceae (Sekereš et al., 2017). An analogous situation, i.e. the presence of pollen-optimized paralogs, has been documented in the case of SEC3, SEC15 and EXO84 exocyst subunits in Arabidopsis, where SEC3a, SEC15a and EXO84a are pollen-specific, while their siblings are active in the sporophyte (Bloch et al., 2016; Synek et al., 2017; Batystová et al., submitted). Analogically to our observation that the artificial EXO70A2 expression in the sporophyte could complement the sporophytic defects of *exo70a1* mutants, SEC15a, the main SEC15 isoform in pollen, could substitute for the function of SEC15b in the sporophyte, although SEC15a and SEC15b share mere 47% sequence identity at the protein level (Batystová et al., submitted). It is tempting to speculate that exocyst complexes with different subunit composition may bind vesicles with distinct secretory cargos and deliver them to special PM subdomains (due to different EXO70-membrane interactions, see Žárský et al., 2009; Sekereš et al., 2017; Kubátová et al. 2019). In contrast, SEC10a and SEC10b represent a very recent duplication without any signs of expression or functional diversification (Vukašinović et al., 2014).

Based on the phenotypic deviations of *exo70a2* mutants, we documented that EXO70A2 participates in pollen maturation, germination, and pollen tube tip growth. Also mutants in two other exocyst subunits, *sec3a* and *sec8*, exhibited severe defects in their germination efficiency (Bloch et al., 2016; Cole et al., 2005). One of the crucial regulatory modules in pollen germination and pollen tube growth is the ROS production (Potocký et al., 2007; Potocký et al., 2012; Smirnova et al., 2014). Two Arabidopsis pollen-specific NADPH oxidases, RBOHH and RBOHJ, localized at the PM are necessary for ROS accumulation in pollen grain cell wall upon pollination and later in pollen tube tips. Double mutants in these genes exhibited much lower ROS level resulting in collapsing pollen tubes (Lassig et al., 2014; Kaya et al., 2015). Interestingly, data of Smirnova et al. (2014) suggest that targeted production of extracellular ROS is required in order to change mechanical properties of intine pollen grain cell wall to ensure germination. Since we observed a significant decrease in ROS production in *exo70a2* germinating pollen grains, we suggest that the pollen exocyst may function in the delivery of NADPH oxidases to specific sites
at the PM during germination.

Following pollen germination, an elongating pollen tube requires precisely regulated secretory machinery to support the intensive tip growth. Mutants in several core exocyst subunits (*sec5a/b, sec6, sec8, sec15a*) generate extremely short and wide pollen tubes with drastically reduced fertilization capacity similarly to *exo70a2* described here (Hála et al., 2008). This pollen tube phenotype is most likely caused by an inefficient targeting of secretory vesicles during the tip growth, because the deposition of major cell wall components was otherwise normal. In contrast, the mutant in *EXO70C2*, a member of the EXO70.2 clade, exhibited very different phenotype of mutant pollen tubes, caused by impaired regulation of growth rate, which pointed to a diverged function of EXO70C2 (and its close paralog EXO70C1) independent of the conventional exocyst function (Synek et al., 2017). Furthermore, EXO70A2 showed the PM localization identical to core exocyst subunits, SEC8, SEC10a and SEC15a in growing pollen tube tips in Arabidopsis (Synek et al., 2017; Batystová et al., submitted). At the molecular level, EXO70A2 probably mediates the exocyst interaction with the PM similarly to Exo70 in yeast and mammalian cells (Boyd et al., 2004; He et al., 2007; Pleskot et al., 2015). This notion is supported by our observation that the N-terminally truncated SEC3a subunit in Arabidopsis pollen was unable to interact with the PM, but still localized to the apical PM in growing pollen tube tips as a part of the complex (Bloch et al., 2016).

Although the transmission efficiency of the *exo70a2* mutant allele was heavily impaired, the mutation did not cause a complete transmission defect like that observed in exocyst loss-of-function mutants in the core exocyst subunits (*sec6, sec8, sec15a*) (Hála et al., 2008). This indicates that some other EXO70 isoform(s) could provide a partially overlapping function to EXO70A2; possible candidates include EXO70H3, H5 and H6. However, their abundance in the pollen proteome is most likely very low (Synek et al., 2017) and their relevance thus would have to be proved experimentally. Alternatively, *EXO70A1* might be activated when *EXO70A2* is disrupted, despite the fact that *EXO70A1* is normally not expressed in pollen (Synek et al., 2006; www.Genevestigator.com – Hruz et al., 2008). In general, we propose that the other six EXO70 isoforms expressed in pollen adopted specific functions with no or only partial functional overlap with EXO70A2. In addition, their activities might be restricted to certain stages of pollen development or specific endomembrane domains.

Proper cell wall deposition is essential for efficient pollen germination and pollen tube growth (Chebli et al., 2012; Leroux et al., 2015; MacAlister et al., 2016). Since the exocyst is responsible for efficient polarized delivery of cell wall materials and cell wall modifying enzymes, it is not surprising that *exo70a2* mutants exhibit defects in both pollen germination and pollen tube growth. Similar phenotypes were reported also for several Arabidopsis mutants in pollen-specific pectin methylesterases (PME). Notably, the homozygous *pme48* mutant showed a significant delay in the pollen grain germination and produced significantly wider pollen tubes (Leroux et al., 2015). Loss of *PPME1* function lead to reduced growth rate and increased width of pollen tubes (Tian et al., 2006). The disruption of PME *VANGUARD1* resulted in impaired pollen tube elongation *in vivo* and bursting of pollen tubes *in vitro* (Jiang et al., 2005). The excessive amount of methylesterified pectins in the intine cell wall of pollen grains and at the pollen tube tip in PME mutants, reduced the Ca^2+^ binding and formation of pectate gel, thus decreasing the rigidity of the cell wall (Leroux et al., 2015). Moreover, higher amount of hydrophobic methylesterified pectins in PME mutants also decreased the rate of pollen grain water uptake during imbibition, affecting the germination efficiency (Leroux et al., 2015). Other cell modifying enzymes are also important for pollen tube tip growth. For example, mutants in hydroxyproline O-arabinosyltransferases, HPAT1 and HPAT3, involved in modification of cell wall-associated extensins, exhibited a similar phenotype to *exo70a2* regarding their defect in pollen tube elongation (MacAlister et al., 2016). Higher percentage of non-viable pollen grains and delayed germination in *exo70a2* could be also explained by mislocalization or inefficient delivery of some cell wall modifying enzymes (or other cell wall components) whose delivery depends on EXO70A2-containing exocyst complex during pollen maturation and germination.

In summary, we conclude that the duplication of *EXO70A* genes to sporophyte- and male gametophyte-specific paralogs occurred independently in monocots and dicots. We proved that among seven EXO70 isoforms expressed in Arabidopsis male gametophyte EXO70A2 is indeed the main conventional EXO70 isoform contributing to the canonical exocyst function in polarized secretion – analogically to its sibling, EXO70A1, in the sporophyte. However, EXO70A2 seems to play several roles in the male gametophyte, because it is important not only for the pollen tube tip growth, but also for the previous phases – pollen grain maturation and germination. Importantly, despite the deep evolutionary split, EXO70A2 still keeps the ability to fully substitute for the EXO70A1 function in the sporophyte. Overall, we characterized a new molecular component critical for pollen function and therefore for efficient plant sexual reproduction.

## MATERIALS AND METHODS

### Phylogenetic and expression analysis

To perform the phylogenetic analysis, a set of EXO70.1 clade sequences from our previous studies (Cvrčková et al., 2012; Rawat et al., 2017) has been updated according to their newest database status and homologs from additional plant species were obtained by BLASTP searches of the GenBank, Phytozome (Goodstein et al., 2012) and GDR (www.rosaceae.org) databases using Arabidopsis EXO70A1 (At5g03540) and EXO70A2 (At5g52340) sequences as a query. The full list of sequences is provided in Supplemental File S1. Protein sequence alignment was constructed using the MAFFT E-INS-I algorithm (Katoh and Standley, 2013) in Jalview software (Waterhouse et al., 2009). Gaps and non-conserved regions were then manually deleted from the alignment, giving the matrix of 56 taxa and 546 positions. Bayesian phylogeny inference was performed using MrBayes software (Ronquist et al., 2012) with a WAG amino acid model, where the analysis was performed in four runs with four chains and 500,000 generations, and trees were sampled every 100 generations. All four runs asymptotically approached the same stationarity after the first 125,000 generations, which were omitted from the final analysis. Maximum-likelihood phylogeny was performed in Phyml software (Guindon et al., 2010) using the LG matrix, γ-corrected for among-site rate variation with four rate site categories plus a category for invariable sites, with all parameters estimated from the data model to build the phylogenetic tree. Bootstrap analysis (500 replicates) was performed to estimate the probability of maximum-likelihood tree topology.

Analysis of the EXO70.1 paralogues expression in male gametophyte and sporophytic tissues was performed using the CoNekT (Proost and Mutwil, 2018) and GEO (Barrett et al., 2013) tools. Additional anther/pollen RNA-seq expression data for sorghum, Arabidopsis, poplar and tobacco were obtained from (Davidson et al., 2012; Loraine et al., 2013; Zhao et al., 2016; Conze et al., 2017), respectively.

### Plant material and growth conditions

The Arabidopsis *exo70a1-2* (SALK_135462; Synek et al., 2006) and *exo70a2-1* (GABI_824D06; Synek et al., 2017) lines were described previously. The *exo70a2-2* line (FLAG_264F01) was obtained from INRA. Genotypes of individual plants were always analyzed by PCR genotyping (for primers – see Supplemental Tab. S1).

Seeds were surface sterilized (70% ethanol for 3 min, 20% commercial bleach for 5 min, rinsed four times with sterile distilled water) and stratified for 2 days at 4°C. Seeds were germinated on vertical 1/2 MS agar plates (one-half-strength Murashige and Skoog medium [Duchefa Biochemie] supplemented with 1% sucrose, vitamin mixture, and 1% plant agar [Duchefa Biochemie]) at 22°C under long-day conditions (16 h light/8 h dark cycles). Seedlings were transferred to turf pellets (Jiffy Products International, Norway) after 8 days and grown until harvest at the same growth conditions.

### Preparation of the *exo70a2* mutant line

The new Arabidopsis mutant line (*exo70a2-3*) was generated using the egg cell-specific promoter-controlled CRISPR/Cas9 technology, employing the pHSE401E vector (Wang et al., 2015). To increase the chance of a mutation, two target single-guide RNAs (sgRNAs) were designed. Flowering Arabidopsis plants of the Col-0 ecotype (NASC collection) were transformed using the *Agrobacterium tumefaciens*-mediated floral dip (Clough and Bent, 1998). After selection on hygromycin, a mutant line was identified by sequencing of PCR products obtained from the targeted part of the *EXO70A2* gene (At5g52340) having a 13bp insertion in the expected site (…TCGAGCTGCGGT<u>ttcatgcgatttt</u>GTTGGAACAGAG…). The genotype was then analyzed by PCR genotyping, when one of each pair of primers for the wild-type and mutant allele, respectively, were designed over the CRISPR-modified site (see Supplemental Fig. S1A and Tab. S1 for primers).

### Semi-quantitative RT-PCR analysis of the *EXO70A2* transcription

To determine the *EXO70A2* transcript level in the CRISPR-modified mutant line, total RNA was extracted from 100 mg of apical parts of primary inflorescences of *exo70a2* and WT plants using the RNeasy kit (Qiagen). Prior to reverse transcription, samples were treated by DNase I (New England Biolabs). The cDNA was synthesized using oligo-dT primers, 2 μg of total RNA and Transcriptor High Fidelity cDNA Synthesis kit (Roche). Transcript abundance was assayed by semi-quantitative PCR for two gene regions – up-stream and down-stream from the CRISPR-generated insertion (see Supplemental Fig. S1) – using *EXO70A2*-specific primers; the primers were especially checked for potential interference with *EXO70A1* and *EXO70A3* paralogs (Supplemental Tab. S1). ACTIN7-specific primers were used as a quantitative control. The optimal number of PCR cycles was determined empirically.

### Analysis of pollen development

Pollen grains at different developmental stages were collected from a series of flower buds ending with a freshly open flower and used for further analyses of nuclear division and pollen grain viability.

To analyze the developmental progress, pollen grains were stained with 4’,6-diamino-2-phenylindole (DAPI) at final concentration 4 μg/ml in PIB buffer (100 mM Na_3_PO_4_, pH = 7.5; 1 mM EDTA; 0.1% [v/v] Triton X-100) as described by Backues et al. (2010). Images were captured using a Zeiss Axio Imager 2 microscope with an EC Plan-Neofluar 40×/0.75 objective, a filter set for observation of blue fluorescence, DIC optics, and Zeiss Axiocam 506 Color camera.

To inspect the pollen grain viability we used Alexander staining (Alexander, 1969) with following modifications: 10× diluted, 15 min incubation time at room temperature. Images were captured using a Nikon Eclipse 90i microscope with Plan Apo 10×/0.45 objective, DIC optics, and a Zyla sCMOS camera (Andor). Fully developed non-viable (not stained) pollen grains, but not collapsed underdeveloped grains, were counted.

### Germination of Arabidopsis pollen *in vitro*

Pollen was germinated on a thin layer of semi-solid pollen germination medium (10% sucrose, 1.5% low-melting-point agarose (Agarose-Universal, peqGOLD, VWR), 0.01% H_3_BO_3_, 5 mM CaCl_2_, 5 mM KCl and 1 mM MgSO_4_) in chambered Lab-Tek II coverglass (Thermo Scientific). Pollen grains from fully opened flowers were spread onto the medium layer, chambers were closed and placed into standard plant growth conditions (see above).

To evaluate the germination efficiency, pollen grains were supposed as germinated when a pollen tube length reached at least one half of pollen grain diameter. For each genotype, 10 samples originating from 4 different plants were observed. Pictures of germinated pollen were taken using Nikon Eclipse 90i microscope with Plan Apo 10×/0.45 objective. Pollen tube lengths were measured in ImageJ and histograms were generated in Microsoft Excel.

### ROS staining

Production of superoxide was determined by its ability to reduce nitroblue tetrazolium (NBT) to formazan precipitate (Rossetti and Bonatti, 2001). Pollen grains were spread on semi-solid pollen germination medium (see above). After 20 min of imbibition it was overlaid with 20μl-drops of NBT diluted in liquid pollen germination medium (final concentration 2 mg/ml), incubated for 10 min, and immediately imaged using a Nikon Eclipse 90i microscope with Plan Apo 20×/0.75 objective, excluding DIC filters, and maximum field aperture opening. NBT signal intensity was measured in ImageJ. Values were normalized to non-stained pollen grains and related to maximal staining intensity value across all samples. The experiment was repeated in three replicas with similar results.

### Pollen tube morphology and growth rate

Details of pollen tube morphology were captured using Zeiss Axio Imager 2 microscope with an EC Plan-Neofluar 40×/0.75 objective 3 or 16 hours after imbibition for wild type and *exo70a2*, respectively. The pollen tube growth rate was analyzed using the same microscope but with EC Plan-Neofluar 20×/0.5 objective. Time-lapses at a 2 min interval were recorded for 30 min (starting 3 h after imbibition) or for 90 min (starting 16 h after imbibition), for WT and *exo70a2*, respectively. Only pollen tubes longer than twice the pollen grain diameter were evaluated, because those already had reached the maximal growth rate.

### Callose visualization in Arabidopsis pistils

Self-pollinated pistils (at least 30 for each genotype) from fully opened flowers were collected and stained with aniline blue according to Mori et al. (2006) and imaged using a Nikon Eclipse 90i microscope with a Plan Apo 4×/0.2 objective and a Zyla sCMOS camera (Andor). Length of the longest pollen tube in every pistil was measured, and recorded values were evaluated statistically using Student’s t-test.

### Fluorescent staining of cell wall components

Propidium iodide, Calcofluor White or Aniline Blue was diluted in liquid germination medium and gently applied onto pollen germinated *in vitro* immediately before imaging. Images were captured using a Zeiss LSM 880 confocal laser scanning microscope with Plan-Apochromat 10×/0.45, Plan-Apochromat 20×/0.8, C-Apochromat 40×/1.2 WI, and C-Apochromat 63×/1.2 WI objectives. Working concentrations, excitation laser wavelengths, and the range of recorded emission wavelengths were as follows: propidium iodide, 30 mM, 514 nm, and 566–719 nm; Calcofluor White, 1 mg/ml, 405 nm, and 410–523 nm; decolorized Aniline Blue, 0.001%, 405 nm, and 410–535 nm.

### Cloning, complementation assays and imaging of GFP-tagged EXO70A2

All constructs were prepared using the MultiSite Gateway^®^ (Invitrogen). For preparation of the *pEXO70A2∷GFP:EXO70A2* construct the EXO70A2 promoter (1,068 bp upstream from the start codon) was subcloned into pENTR 5’-TOPO; the GFP gene in pEN-L1-F-L2 was obtained from VIB in Ghent (Karimi et al., 2007); the *EXO70A2* (At5g52340) with the stop codon was subcloned into pDONR P2R-P3 using the Gateway BP clonase (Invitrogen). The three elements were then recombined together into the pB7m34GW destination vector (Karimi et al., 2007) using the Gateway LR clonase (Invitrogen).

For cloning of *pEXO70A1∷GFP:EXO70A2* and *pEXO70A1∷GFP:EXO70A1*, the *EXO70A1* promoter (1kb upstream from the start codon) was subcloned into pDONR P4-P1r. The *EXO70A1* was subcloned into pDONR P2R-P3. Then, multi-site reactions were performed to assemble the *EXO70A1* promoter, GFP, and *EXO70A1* or *EXO70A2* into the destination vector pB7m34GW. The insertions in pDONR vectors were sequenced using M13 primers. The final constructs were sequenced using M13 primers and two GFP primers (see Supplemental Tab. S1).

The final *pEXO70A2∷GFP:EXO70A2* construct was then introduced to *exo70a2* heterozygous plants, while *pEXO70A1∷GFP:EXO70A1* and *pEXO70A1∷GFP:EXO70A2* to *exo70a1* heterozygous plants, using the *Agrobacterium tumefaciens*-mediated floral dip method (Clough and Bent, 1998). Transformed seedlings were selected on BASTA. Complementation assays were performed using 3 or more independent transformed lines for each construct.

The subcellular localization of GFP:EXO70A2 in pollen grains, pollen tubes and roots was performed using a Zeiss LSM 880 confocal laser scanning microscope equipped with C-Apochromat 63×/1.2 WI. The fluorophore was excitation with a 488-nm laser, and emitted fluorescence was recorded at 493–535 nm. The pollen was germinated as described above and carefully transferred to the microscope 2 h after imbibition.

## ABBREVIATIONS

CDS: coding sequence
GFP: green fluorescent protein
NBT: nitroblue tetrazolium
PM: plasma membrane
PME: pectin methylesterases
ROS: reactive oxygen species
WT: wild type

## SUPPLEMENTAL DATA

**File S1.** EXO70 sequences used for the phylogenetic analysis.

**Table S1.** List of primers used in this study.

**Figure S1.** Characterization of the *exo70a2* mutant line generated using the CRISPR/Cas9 system.

**Figure S2.** Development of WT and *exo70a2* pollen.

**Figure S3.** Pollen germination efficiency is normal in complemented *exo70a2* mutant plants.

## ACKNOWLEDGEMENTS

This work was supported by the Czech Science Foundation (GAČR) – project 18-18290J, and part of the
V.Z. income was covered by the Ministry of Education, Youth and Sports of the Czech Republic from European Regional Development Fund-Project “Centre for Experimental Plant Biology” CZ.02.1.01/0.0/0.0/16_019/0000738.

Microscopy was performed in the Laboratory of Confocal and Fluorescence Microscopy and the Microscopic facility of IEB. These facilities were supported by the European Regional Development Fund-Project and the state budget of the Czech Republic (projects no. CZ.1.05/4.1.00/16.0347 and CZ.2.16/3.1.00/21515), the Operational Programe Prague – Competitiveness (project no. CZ.2.16/3.1.00/21519), and the Czech-BioImaging large RI project LM2015062.

We thank to Marta Čadyová for technical support and Juraj Sekereš for critical reading of the manuscript.

