## Supplemental Table and Figures for "EXO70A2 is critical for the exocyst complex function in Arabidopsis pollen"

**Table S1.** List of primers used in this study.

| Primer name | Primer sequence | Use |
|---|---|---|
| *Preparation of the exo70a2 CRISPR line* | | |
| EXO70A2 DT1-BsF | ATATATGGTCTCGATTGGATACTCGAGCTGCGGTGTGTT | assembly of two gRNA expression cassettes in pHSE401E vector |
| EXO70A2 DT1-F0 | TGGATACTCGAGCTGCGGTGTGTTTTAGAGCTAGAAATAGC | |
| EXO70A2 DT2-R0 | AACGCGGAAGCTAAAATATTGAGAATCTCTTAGTCGACTCTAC | |
| EXO70A2 DT2-BsR | ATTATTGGTCTCGAAACGCGGAAGCTAAAATATTGAGAA | |
| *Plant genotyping* | | |
| gtyp-wt-FW | GCAGCTGAGGTTATATTGGATCAATTTG | gtyp-wt FW+RV detects WT allele |
| gtyp-wt-RV | AAACTCTGTTCCAACACCGCAG | |
| gtyp-crispr-FW | ATCAAAGCTAGAAGATGAGTTCAGACA | gtyp-crispr FW+RV detects CRISPR insertion |
| gtyp-crispr-RV | CTGTTCCAACAAAATCGCATGAAAC | |
| exo70a1-2 LP | TCCATGGACACAAATTTCATG | LP + RP detects WT allele |
| exo70a1-2 RP | TCTACTGGCATTTTCCCAATG | RP + LBb1.3 detects T-DNA |
| LBb1.3 | ATTTTGCCGATTTCGGAAC | |
| *Cloning of EXO70A2 and EXO70A1* | | |
| A2_prom_FW | TCTAAAGGATGATTTTTTCGTCA | A2_prom_FW + A2_prom_RV used for cloning of the EXO70A2 promoter |
| A2_prom_RV | TTTCTTGATTTGGATCGATGAAC | |
| A2_FW | GGGGACAGCTTTCTTGTACAAAGTGGCTATGGGGGTGGCTC | A2_FW (attB2r) + A2_RV(attb3) used for |

| | | |
|---|---|---|
| A2_RV | GGGGACAACTTTGTATAATAAAGTTGCTTTATCTCTTTGGCTCACTCC | cloning of the EXO70A2 gene |
| A1_prom_fw | ATTGTAAAAAGGGAATGAGCAT | A1_prom_fw (attB4) + A1_prom_rev (attB1R) used for cloning of the EXO70A1 promoter |
| A1_prom_rev | AAAATAACGAATAATCTTTCTGAGTTGA | |
| A1_FW | CTTGTACAAAGTGGCTATGGCTGTTGATAGCAGA | A1_FW (attB2r) + A1_RV (attB3) used for cloning of EXO70A1 gene |
| A1_RV | GTATAATAAAGTTGTTACCGGCGTGGTTC | |
| attB2r adaptor | GGGGACAGCTTTCTTGTACAAAGTGG | |
| attB3 adaptor | GGGGACAACTTTGTATAATAAAGTTG | |
| attB1R adaptor | GGGGACTGCTTTTTTGTACAAACTTG | |
| attB4 adaptor | GGGGACAACTTTGTATAGAAAAGTTGAA | |

*Sequencing of EXO70A2 constructs in Gateway vectors*

| | | |
|---|---|---|
| M13_FW | GTAAAACGACGGCCAGT | |
| M13_RV | AACAGCTATGACCAT | |
| GFP_seq_FW | CCACAACGTCTATATCATGG | |
| GFP_seq_RV | ACGCCGTAGGTCAG | |
| A2_CRISPR_FW | GGACCCAATCCAGTTAGATATCAAGTA | |
| A2_CRISPR_RV | GGCAACAAAAGCAAGTGCC | |
| *RT-PCR analysis* | | |
| A2_5'-FW | TGGCTCAAGCAATGGAAGCCCTAA | A2_5'-FW + A2_upstream-RV amplify 5' half of EXO70A2 upstream of the CRISPR insertion |
| A2_upstream-RV | GAGGAATGACAGTTGGGACAGTAAAAATA | |
| A2_downstream-FW | TGAGGTCACTGTAAATAGTGTAGCTG | A2_downstream-FW + A2_3'-RV amplify 5' half of EXO70A2 upstream of the CRISPR insertion |
| A2_3'-RV | TCTCTTTGGCTCACTCCATGTCTTG | |

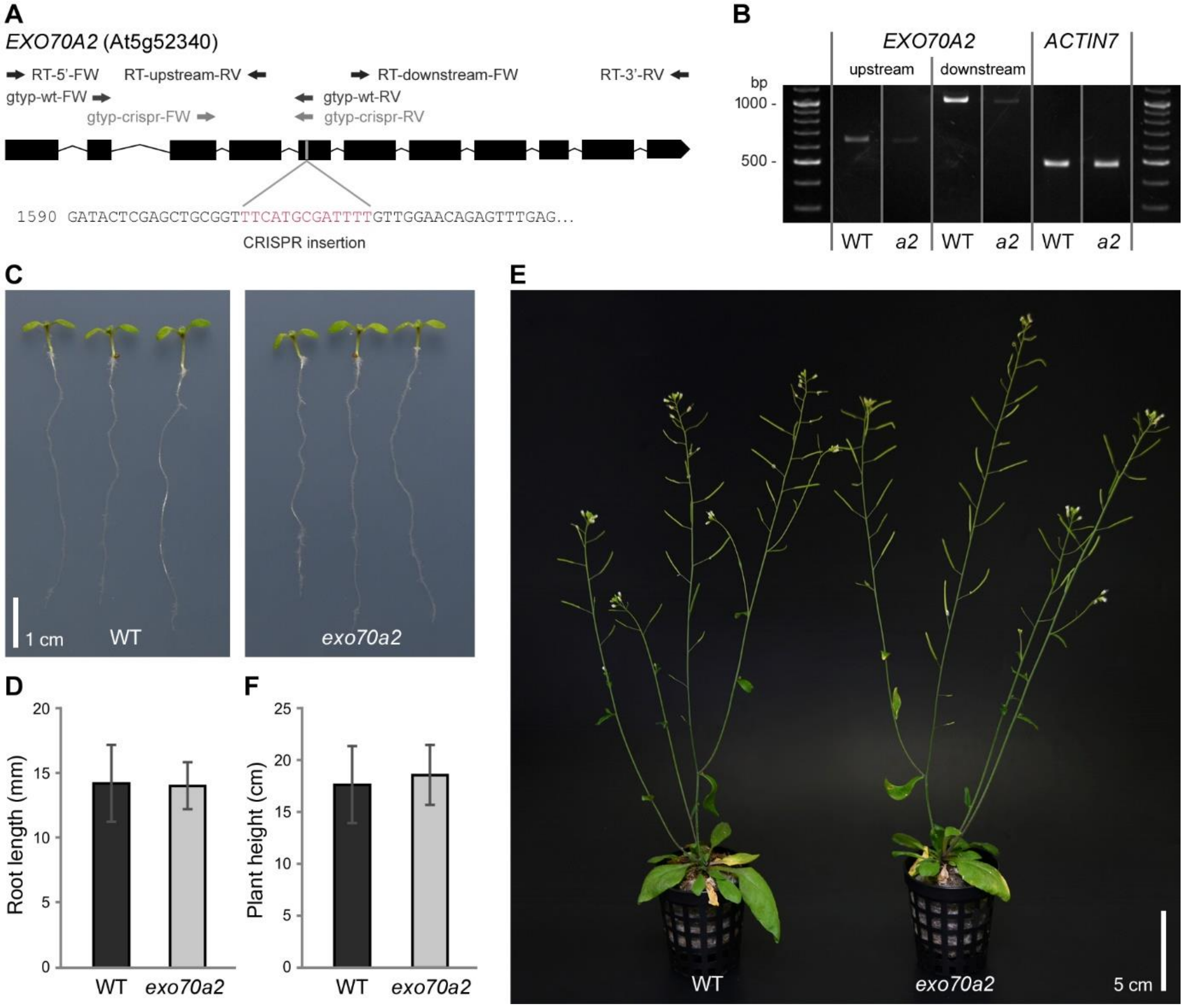


**Figure S1.** Characterization of the *exo70a2* mutant line generated using the CRISPR/Cas9 system.

A) Schematic display of the *EXO70A2* gene with the CRISPR-generated insertion. Position of primers used for genotype and RT-PCR analyses are indicated (see Table S1).
B) Semi-quantitative RT-PCR analysis on cDNA samples prepared from WT and *exo70a2* inflorescences. The *EXO70A2* transcript level was inspected in two gene regions, upstream and downstream from the CRISPR-generated insertion, using pairs of primers indicated in A. Housekeeping *ACTIN7* was analyzed as a quantitative control.
C) Representative 7-day-old WT and homozygous *exo70a2* seedlings.
D) Primary root lengths of 7-day-old seedlings are not significantly different (n > 101; t-test p_value = 0.541).
E) Representative 40-day-old WT and homozygous *exo70a2* plants.
F) Plant heights of 40-day-old plants are not significantly different (Student's t-test p_value = 0.528).

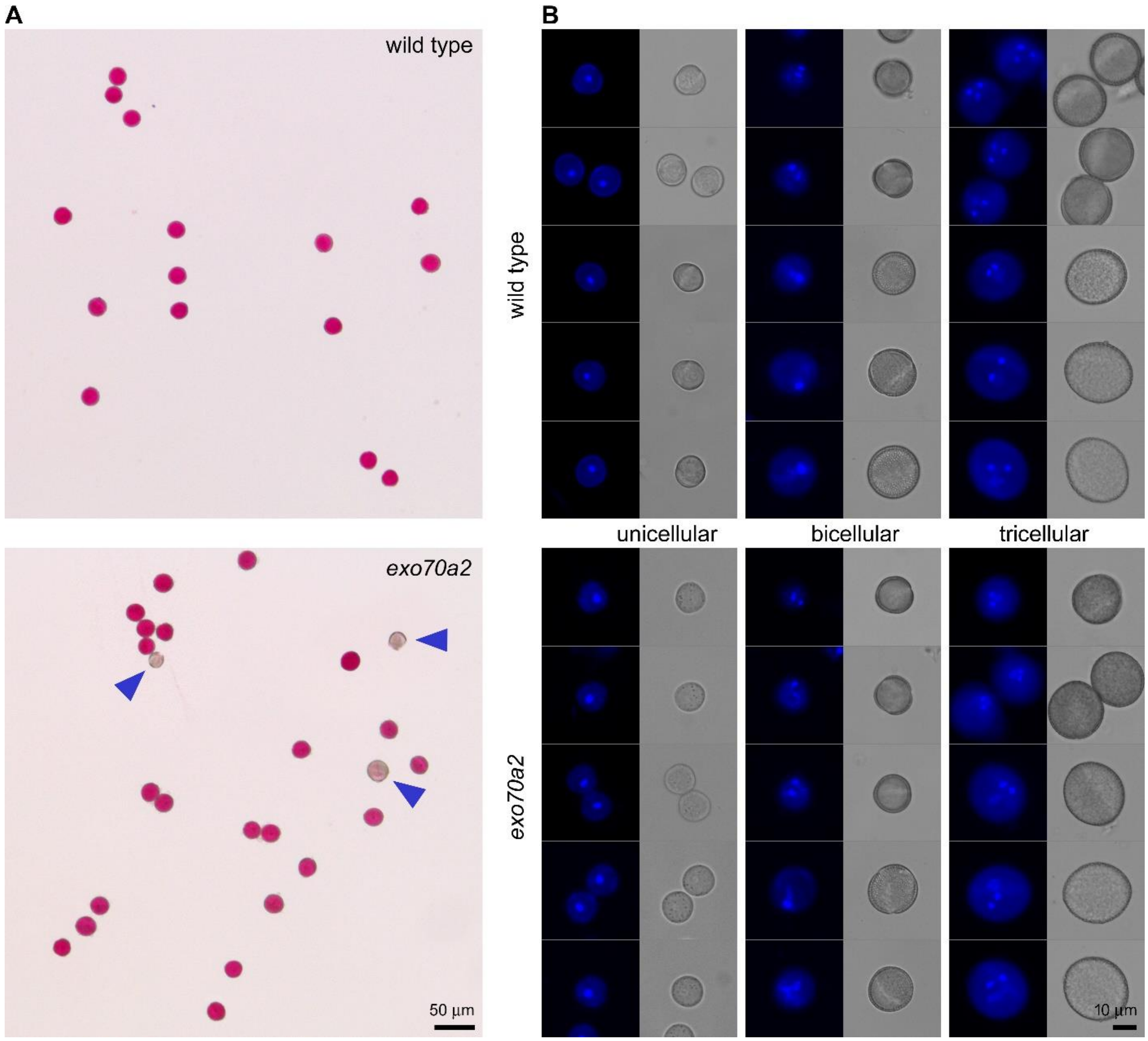


**Figure S2.** Development of WT and *exo70a2* pollen.

A) Viability of pollen grains visualized by Alexander staining (viable grains are in purple, non-viable are pale – marked by arrowheads).
B) Stages of pollen development stained by DAPI and complemented by brightfield images.

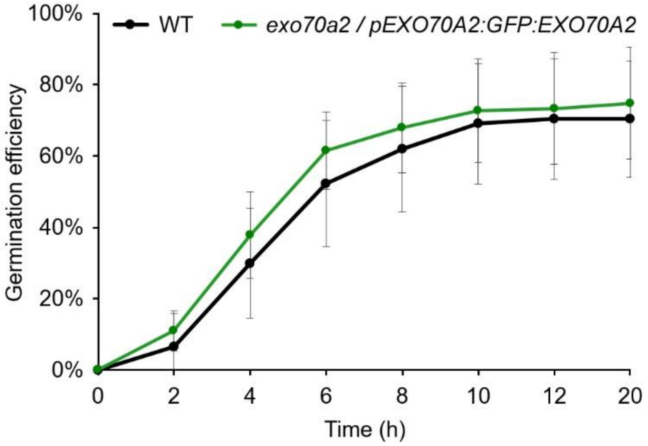


**Figure S3.** Pollen germination efficiency is normal in complemented *exo70a2* mutant plants.

Pollen germination efficiency of *exo70a2* and WT pollen *in vitro* at different time points.
